# Retrieving the near-complete genome of a threatened bird from wild frozen samples

**DOI:** 10.1101/2022.12.22.521577

**Authors:** Haoran Luo, Xinrui Jiang, Boping Li, Jiahong Wu, Jiexin Shen, Zaoxu Xu, Xiaoping Zhou, Zhen Huang, Minghao Hou, Xiaobin Ou, Luohao Xu

## Abstract

Conservation genomics often relies on non-invasive methods to obtain DNA fragments which limit the power of multi-omic analyses for threatened species. Collecting samples from frozen dead animals in the wild provides an alternative approach to obtaining high-quality nucleic acids. Here, we report multi-omic analyses based on a well-preserved great bustard individual (*Otis tarda*, Otidiformes) of a recent death found in the mountainous region in Gansu, China. We generated a near-complete genome assembly (OTswu) having only 18 gaps scattering in 8 out of the 40 assembled chromosomes. Unlikely most other bird genome assemblies, OTswu contains complete chromosome models (2n = 80). We demonstrated that the great bustard genome likely retained the ancestral avian karyotype. We also characterized the DNA methylation landscapes of OTswu which are strongly correlated with GC content and gene expression. Our phylogenomic analysis suggested Otidiformes and Musophagiformes were sister groups that diverged from each other 46.3 million years ago. The genetic diversity of great bustard was found the lowest among the four available Otidiformes genomes, possibly due to population declines during past glacial periods. As one of the heaviest migratory birds, great bustard possesses several expanded gene families related to cardiac contraction, actin contraction, calcium ion signaling transduction, as well as positively selected genes enriching for metabolism. Finally, we identified an extremely young evolutionary stratum on the sex chromosome, a rare case among Neoaves. Together, our study combining long-read sequencing and RNA-seq technology provides a working strategy for conducting multi-omic analyses for threatened species by retrieving high-quality nucleic acids from dead animals frozen in the wild.

## 1 Introduction

Conservation genetics has moved towards an era where high-quality reference genomes are often required (Lee et al., 2016; Formenti et al., 2022). For threatened large animals, one of major challenges in conservation genomics is to obtain fresh samples for genome sequencing, in particular long-read sequencing (Irestedt et al., 2022). Noninvasive sampling (Pierre Taberlet, Waits, & Luikart, 1999), including collecting hairs, feathers (Higuchi, von Beroldingen, Sensabaugh, & Erlich, 1988; Morin, Wallis, Moore, Chakraborty, & Woodruff, 1993), feces(Hoss, Kohn, Paabo, Knauer, & Schroder, 1992), or museum specimens (Feng et al., 2019) has been widely used in conservation biology, but severe RNA degradation, highly fragmented DNA and heavy contamination limit the performance of high-quality DNA extraction or transcriptome profiling (P. Taberlet, Mattock, Dubois-Paganon, & Bouvet, 1993; Pierre Taberlet & Luikart, 1999). The degeneration of DNA and RNA is much slower in cold temperatures, therefore sampling animals of a recent death in frigid zones provides an alternative strategy for obtaining well-preserved DNA or RNA. Retrieving well-preserved DNA from wild frozen samples for genome assembly and population genetic analyses has been successful for several mammalian species(S. Liu et al., 2014; Palkopoulou et al., 2015), including the extinct mammoth (Palkopoulou et al., 2015; Rogers & Slatkin, 2017; van der Valk et al., 2021), but such study for avian species has been rare (Irestedt et al., 2022).

The recent development of long-read sequencing provides an unprecedented opportunity for complete genome assembly, or telomere-to-telomere (T2T) genome assembly (Nurk et al., 2022). This is a critical step towards building a “Digital Noah’s Ark” as a resort to preserve genomic information for endangered species (Wei et al., 2022). Nanopore technology provides a fast and high throughput method for sequencing long-reads that can be up to hundreds of kilobases and has been widely used in conservation genomics (Watsa, Erkenswick, Pomerantz, & Prost, 2020). In addition, Nanopore reads contain DNA methylation signals, allowing for the identification of genome-wide epigenetic modifications (Laszlo et al., 2013; Schreiber et al., 2013) that are critical for the maintenance of genome stability and gene expression regulation.

Birds have the most streamlined genomes among vertebrates where large-scale genome sequencing projects have been overwhelmingly successful (G. Zhang et al., 2014; Feng et al., 2020). To date, over 500 bird genomes are available, though most were sequenced with short reads (Bravo, Schmitt, & Edwards, 2021). A recent chicken pan-genome study using long-read sequencing, however, suggests bird genomes are far from complete, missing thousands of genes previously thought to be lost (M. Li et al., 2022). The Vertebrate Genome Project recently also reports massive false gene losses in bird genomes (J. Kim et al., 2022). Moreover, a few microchromosomes have been missing or incomplete in bird genome assemblies likely due to their high GC content and accumulation of simple repeats (Peona et al., 2021; J. Kim et al., 2022).

The great bustard (*Otis tarda*) has been a vulnerable species according to the IUCN Red List of Threatened Species since 1988. Its habitat distributes across the Eurasian continent and was thought to be extinct in England. The decline of great bustard population in recent years is mainly caused by habitat loss due to human activity (Bankovics, 2003), collisions with power lines (Raab, SchÜTz, Spakovszky, Julius, & Schulze, 2011) and hunting. Great bustard is amongst the heaviest living flying animals. The male can range in weight from 5.8 to 18 kg (del Hoyo, Elliot, & Sargatal, 1996). The heaviest verified specimen was about 21 kg, a world record for the heaviest flying bird (Wood & Gerald, 1982). The great bustard is also one of the most sexually dimorphic birds in body size with adult male great bustards being ~2.5 times heavier than females (Alonso et al., 2009). Despite the heavy body size, great bustard is a powered flier and can reach speeds of 48 km/h to 98 km/h during migration (Kessler, Batbayar, Natsagdorj, Batsuur’, & Smith, 2013), and can migrate over 2,000km in northern Mongolia breeding populations (Kessler et al., 2013). Great Bustards are normally less active in snowy and frosty conditions (Litzbarski, Litzbarski, & Petrick, 1987), likely due to the need to conserve energy (Streich, Litzbarski, Ludwig, & Ludwig, 2005). During the snowy night, great Bustards often prefer to remain motionless in one place, causing the snow under them to melt (Streich et al., 2005).

In the January of 2022, we spotted a dead adult great bustard in a mountainous region of Gansu, China. The cause of death was unidentified, and the date of death was unclear. We immediately relocated the animal to the lab for dissection, and extracted samples for Nanopore ultra-long, Hi-C and RNA-seq library preparation. Fortunately, both DNA and RNA was well preserved, and we were able to retrieve a near-complete genome assembly that contained complete chromosome models. Combining genomic, epigenomic and transcriptomic analyses, we illustrated the genetic diversity, demography, gene expression landscape and the evolutionary history of the sex chromosomes of great bustard. Overall, our study provides a working strategy for retrieving the near-complete genome from an individual of a recent death in the wild, enabling epigenetic and transcriptomic analyses for endangered species.

## 2 MATERIALS AND METHODS

### 2.1 Sample collection and sequencing

An adult male great bustard was found dead on the 17th of January, 2022, at Hesheng Town, Ning County, Qingyang City, Gansu Province, China (35°431255’N, 107°782055’E) (**Supplementary Fig. S1**). The local temperature was −7 to 4 °C. The frozen individual was immediately relocated to the laboratory for dissection. Five tissues were collected for sequencing, including leg thigh muscle, brain, heart, lung, and liver.

Genomic DNA for ultra-long reads sequencing was isolated from the muscle. Ultra-long DNA was extracted by the SDS method without a purification step to ensure a sustained length of genomic DNA. Then, length >50 kb genomic DNA was size-selected in the SageHLS HMW library system (Sage Science, USA), and sequencing libraries were processed by using the Ligation sequencing 1D kit (SQK-LSK109, Oxford Nanopore Technologies, UK) according to the manufacturer’s instructions. Three DNA libraries were constructed and sequenced on the PromethlON (Oxford Nanopore Technologies, UK) at the Genome Center of Grandomics (Wuhan, China).

For short-read sequencing, the cetyltrimethylammonium bromide (CTAB) method was used to extract genomic DNA from thigh muscle. Genomic DNA was randomly fragmented, then an average size of 200-400 bp fragments was selected by Agencourt AMPure XP-Medium kit. The constructed library was then sequenced on the MGISEQ-T7 platform with PE 150 bp mode.

Hi-C sequence was used to anchor assembled contigs onto the chromosome. Muscle tissues were cut into pieces in nuclei isolation buffer supplemented with 2% formaldehyde for cross-linking. Cross-linking was stopped by adding glycine and additional vacuum infiltration. Fixed tissue was then grounded to powder and resuspended in nuclei isolation buffer to obtain a suspension of nuclei. To remove unligated DNA fragments, the purified nuclei were digested with 100 units of Dpnll and marked by incubating with biotin-14-dATP. Non-ligated DNA was removed owing to the exonuclease activity of T4 DNA polymerase. The ligated DNA was sheared into 300-600 bp fragments, and then followed by a standard library preparation protocol (Meyer & Kircher, 2010). Hi-C sequencing was conducted on the MGISEQ-T7 platform with the PE 150 bp mode.

### 2.2 Genome assembly and assessment

We used three cells of Nanopore ultra-long reads for de novo assembly with default parameters in Nextdenovo v2.5.0 (GrandOmics, 2022). To correct assembly errors, we applied two rounds of polishing with Nextpolish v1.4.0 (Hu, Fan, Sun, & Liu, 2020) based on the long-read alignments. We further polished the assembly twice with short reads using Pilon v1.24 (Walker et al., 2014). Benchmarking Universal Single-Copy Orthologues (BUSCO V5.2.2) (Manni, Berkeley, Seppey, Simao, & Zdobnov, 2021) with the aves_odb10 lineage (n = 8338) was used to assess genome assembly completeness.

### 2.3 Chromosome-level assembly with Hi-C data

We mapped the reads from the Hi-C library sequencing against the contigs with the Juicer (v1.6) pipeline (Durand, Shamim, et al., 2016). The “.hic” file was generated using the 3D-DNA (v180419) pipeline (Dudchenko et al., 2017) and the Hi-C heatmap was visualized in the Juicebox Assembly Tools (Durand, Robinson, et al., 2016) for manual curations.

### 2.4 RNA-sequencing and transcriptome assembly

Total RNA from Leg thigh muscle, brain, heart, lung, and liver tissues were extracted using the Trizol (Invitrogen, Carlsbad, CA, USA) method. RNA-sequencing libraries were prepared and sequenced on the MGISEQ-T7 platform with the PE 150 bp mode. After trimming RNA-seq reads with Trimmomatic (v0.39, default parameters) (Bolger, Lohse, & Usadel, 2014), we mapped the filtered RNA-seq against the genome assembly with HISAT2 (v2.1.0) (D. Kim, Langmead, & Salzberg, 2015). Transcripts were assembled using StringTie (v2.0) (Kovaka et al., 2019). TransDecode (v5.5.0) (Haas et al., 2013) was used to predict protein-coding regions of the assembled transcripts.

### 2.5 Repeat annotation

Avian homology repetitive elements were acquired from RepeatMasker (http://www.repeatmasker.org) database (RepeatMaskerLib.h5). EDTA (v2.0.1) (Ou et al., 2019), TRF (v4.09) (Benson, 1999), and RepeatModeler (v2.0.1) (http://www.repeatmasker.org) were used for *de novo* prediction of repetitive elements. Tandem repeats predicted by TRF (Benson, 1999) were filtered by the pyTanFinder pipiline (Kirov, Gilyok, Knyazev, & Fesenko, 2018). CD-hit (v4.8.1) (W. Li & Godzik, 2006) was used to construct a non-redundant repeat library based on all *de novo* and known libraries. RepeatMasker (v4.1.2-p1) was used to mask repetitive elements in assembled chromosome-level genome with default parameters.

### 2.6 Genome annotation

*Ab initio*, transcriptome-based, and homolog-based approaches were combined to predict protein-coding genes. For homolog-based methods, genome threader (v1.7.1) (Gremme, Brendel, Sparks, & Kurtz, 2005) and exonerate (v2.4.0) (Slater & Birney, 2005) (implement by maker3 (Cantarel et al., 2008)) were used predict gene models. Chicken and zebra finch protein sequences were used as queries for homolog search. For *ab initio*-based methods, AUGUSTUS v3.4.0 (Stanke, Diekhans, Baertsch, & Haussler, 2008), GeneID v 1.4 (Blanco, Parra, & Guigo, 2007) and SNAP (Korf, 2004) were used to predict gene models. The training models of AUGUSTUS were directly acquired based on BUSCO (V5.2.2) gene models. The training set for GeneID was derived from transcriptome-based evidence. The initial training set of SNAP (v2006728) was acquired from homolog proteins and genome-guild transcripts assembled with TRINITY v2.8.5 (Grabherr et al., 2011). Three-round training was performed in SNAP. We then used EVM v1.1.1 (Haas et al., 2013) to integrate all predicted gene models by the above three methods. Finally, we used the PASA (v2.5.2) (Haas et al., 2003) pipeline to polish gene models using TRINITY genome-guild transcript assembly.

Functional annotations were performed with InterProScan (5.35-74.0) (Jones et al., 2014), EggNog-mapper (Huerta-Cepas et al., 2019), and UniProtKB (Boutet, Lieberherr, Tognolli, Schneider, & Bairoch, 2007). BLASTP (Altschul, Gish, Miller, Myers, & Lipman, 1990) was used to search the UniProtKB database with an E-value cutoff of 1E-5. EggNog-mapper v2.1.2 with hmmer (E-value 1e-5) was used to search EggNog v5 Aves (8782) database (Cantalapiedra, Hernandez-Plaza, Letunic, Bork, & Huerta-Cepas, 2021). InterPro databases included COILS, CDD, HAMAP, Gene3D, MobiDB Lite, Panther, Pfam, PIRSF, PRINTS, Prosite, SFLD, SMART, SUPERFAMILY and TIGRfams.

### 2.7 DNA methylation

DNA 5mC methylation was called with Nanopolish (v1.4.0) (Simpson et al., 2017) by using the Hidden Markov Model (HMM). ONT ultra-long fast5 files were used as the input files. The methylation frequency (MF) was calculated as the number of reads on methylated cytosine (xmCpG) divided by the total number of reads covering each cytosine site in the reference (xmCpG + xCpG). We further calculated the mean methylation levels over 50 kb windows, and analyzed the correlations with other genomic features.

### 2.8 Phylogenetic analyses

Coding sequences of chicken (*Gallus gallus*, ASM2420605v1), zebra finch (*Taeniopygia guttata*, GCA_003957565.4), kori bustard (*Ardeotis kori*, ASM1339637v1), Common Cuckoo (*Cuculus canorus*, ASM70932v1), Rock pigeon (*Columba livia*, Cliv_2.1), grey go-away-bird (*Corythaixoides concolor*, ASM1339949v1) and the great bustard (this study) were used to constructed phylogenetic tree and further comparative genomics analyses. To do so, we used OrthoFinder2 (Emms & Kelly, 2015) to identify single-copy orthologous genes for the sampled bird species. IQTREE2 (Minh et al., 2020) was used to construct the reference maximum likelihood tree based on single-copy orthologous genes, and the best substitution model JTT+F+I+I+R4 automatically selected by ModelFinder (Kalyaanamoorthy, Minh, Wong, von Haeseler, & Jermiin, 2017) for protein alignments. For coding sequence (CDS) alignment, the best substitution model GTR+F+R3 was automatically selected by ModelFinder. We used LAST (v1066) (Harris, 2007) to perform whole-genome pairwise alignment with great bustard as the reference, and used MULTIZ (v11.2) (Blanchette et al., 2004) to generate multiple whole-genome alignments. Only one-to-one alignments were retained, and the alignments were further filtered by the TrimAl (v1.2) (Capella-Gutierrez, Silla-Martinez, & Gabaldon, 2009) strict model. The best substitution model GTR+F+R5 was used.

The approximate likelihood calculation method (dos Reis & Yang, 2011) in PAML-MCMCTREE (PAML v4.9j) (Yang, 2007) was used for divergence time estimation. We used PAML-CodeML (Yang, 2007) to acquire an initial branch length with the gradient and Hessian information for the ultrametric tree. Then, this file was taken as the input for MCMCTREE (Yang, 2007) to run 20 million steps of MCMC chains to estimate divergence time. The fossil calibration confidence interval of Neognath and Neoaves (60.2 −86.8 MYA) was derived from the Palaeontologia Electronica Fossil Calibration Database (Benton et al., 2015).

### 2.9 Gene family evolution

The orthogroups identified by OrthoFinder2 were taken by Café (v4.2.1) (De Bie, Cristianini, Demuth, & Hahn, 2006) to infer gene family expansions and contractions. The ultrametric tree topology and branch lengths from the MCMCTREE were used to infer the significance of changes in gene family size in each branch, and significant levels of expansion and contraction (P value) were determined at 0.05. We conducted GO (Gene Ontology) enrichment analysis for genes in the expanded families with PANTHER17.0 (Mi et al., 2021).

### 2.10 Positive selection

We estimated the nonsynonymous to synonymous substitution rate ratios (ω = dN/dS) to assess selection on single-copy orthologous genes. The single-copy orthologous nucleotide sequences were generated by ParaAT 2.0 (Z. Zhang et al., 2012) which generated back-translated nucleotide alignments guided by protein-coding sequences alignments to ensure the alignments were reliable and accurate. MAFFT (v7.505) (Katoh & Standley, 2013) was used as the aligner. All gaps were automatically removed by ParaAT, and alignment lengths shorter than 99 bp were removed. We estimated ω by using the branch-site model (Yang & Nielsen, 2002; J. Zhang, Nielsen, & Yang, 2005) implemented in the CodeML program in PAML (Yang, 2007), using unrooted trees to fit the parameter clock=0. The branch-site model estimated ω values for each site in foreground and background, and then divided sites into three categories: ω□<□ 1 (ω0), ω□=□ 1 (ω1), and ω□>□1(ω2). The null model of the branch-site model does not allow ω larger than 1 in all branches. The alternative hypothesis allowed ω values to be larger than one in the foreground branch, representing the positively selected sites. P-values were calculated through the likelihood ratio test and then adjusted by false discovery rate (FDR) corrections (Benjamini, Krieger, & Yekutieli, 2006). Genes with ω2 higher than 1, FDR smaller than 0.01 and positively selected sites (2a and 2b) more than 5% were considered as positively selected genes (PSGs). We further performed additional two rounds of branch-site model analyses. PSGs missing in any verification round were filtered. Candidate PSGs were taken for GO enrichment analyses in PANTHER17.0 (Mi et al., 2021)

### 2.11 Demographic analyses

We used interval parameters “4+30*2+4+6” sets in PSMC (v0.6.5) (H. Li & Durbin, 2011) which has been used for several bird genomes (Nadachowska-Brzyska, Li, Smeds, Zhang, & Ellegren, 2015; Martini, Dussex, Robertson, Gemmell, & Knapp, 2021; Robledo-Ruiz et al., 2022). We performed 100 times bootstraps to estimate the variance of the simulated results. The neutral mutation rate μ (mutations per base per generation) used for PSMC was μ= D*g / 2T, where D and T were the estimated genome-wide nucleotide divergence and the estimated divergence time between the great bustard and kori bustard, and g was the estimated generation time. Females and males of great bustard have different maturation ages, with males usually starting to mate from 5 to 6 years of age, while females at 2 to 3 years old (del Hoyo et al., 1996). We thus assumed a generation time of 3 years. The genome-wide nucleotide divergence was calculated by nucmer and dnadiff from the MUMmer program (v4.0.0) (Delcher, Phillippy, Carlton, & Salzberg, 2002; Kurtz et al., 2004).

### 2.12 Genetic diversity

Genetic diversity was assessed by calculating individual heterozygosity. Short reads were mapped to genome sequence by using the BWA-MEM algorithm (v0.7.17) (H. Li & Durbin, 2009). Picard package (v2.25.0) (Broad-Institute, 2019) was used to mark duplicates. The Genome Analysis Toolkit (GATK4.2.0.0) (McKenna et al., 2010) HaplotypeCaller module was used for variant calling. SNPs genotypes were generated from GATK GenotypeGVCF s module. Variant quality information Quality by depth (QD) < 2.0, Fisher Strand (FS) > 60.0, RMS Mapping Quality (MQ) < 40.0, Mapping Quality Rank sum (MQRankSum) <-12.5, and Position Rank Sum (ReadPosRankSum) <-8.0 were used to filter low-quality variants by using the VariantFiltration module. Heterozygosity was calculated by dividing the total number of heterozygous sites by the genome size covered by reads.

### 2.13 Sex chromosome evolution

We map resequencing reads from a female individual to the reference genome using BWA MEM. We then used Samtools (v1.9) (H. Li et al., 2009) depth to calculate sequencing converge with default parameters. We also calculated coverage for alignments with a stringent filtering criterion, *i.e*. only retaining reads having no more than 2 mismatches, to avoid ZW cross-mapping. GATK was used to calculate heterozygosity with the same pipeline described above. We used Bedtools (v2.29.1) (Quinlan & Hall, 2010) to calculate the mean converge and heterozygosity over 50 kb windows

## 3 Results

### 3.1 Near-complete genome assembly

We extracted high-molecular-weight DNA from the muscle tissues of the frozen great bustard, and produced 134.1 Gb (~112X genome coverage, **Supplementary Table S1**) Nanopore ultra-long reads. The N50 of the ultra-long reads reached 37.7 kb, suggesting that long fragments of DNA had been preserved. Those ultra-long reads led to a primary assembly with only 129 contigs. The total assembly size is 1.20 Gb, larger than the short-reads-based estimation (1.09 Gb) by 112.6 Mb. The contig N50 is 41.0 Mb, ranking the fourth longest among more than 800 bird genome assemblies available in NCBI, only next to a chicken and two parrots. To correct sequence errors, we polished the contigs using 73.3 Gb short-reads generated based on the same sample (**Fig. 1a**). The BUSCO completeness score is 97.5%, suggesting a high level of assembly completeness (**Supplementary Table S2**).

**Fig 1.**
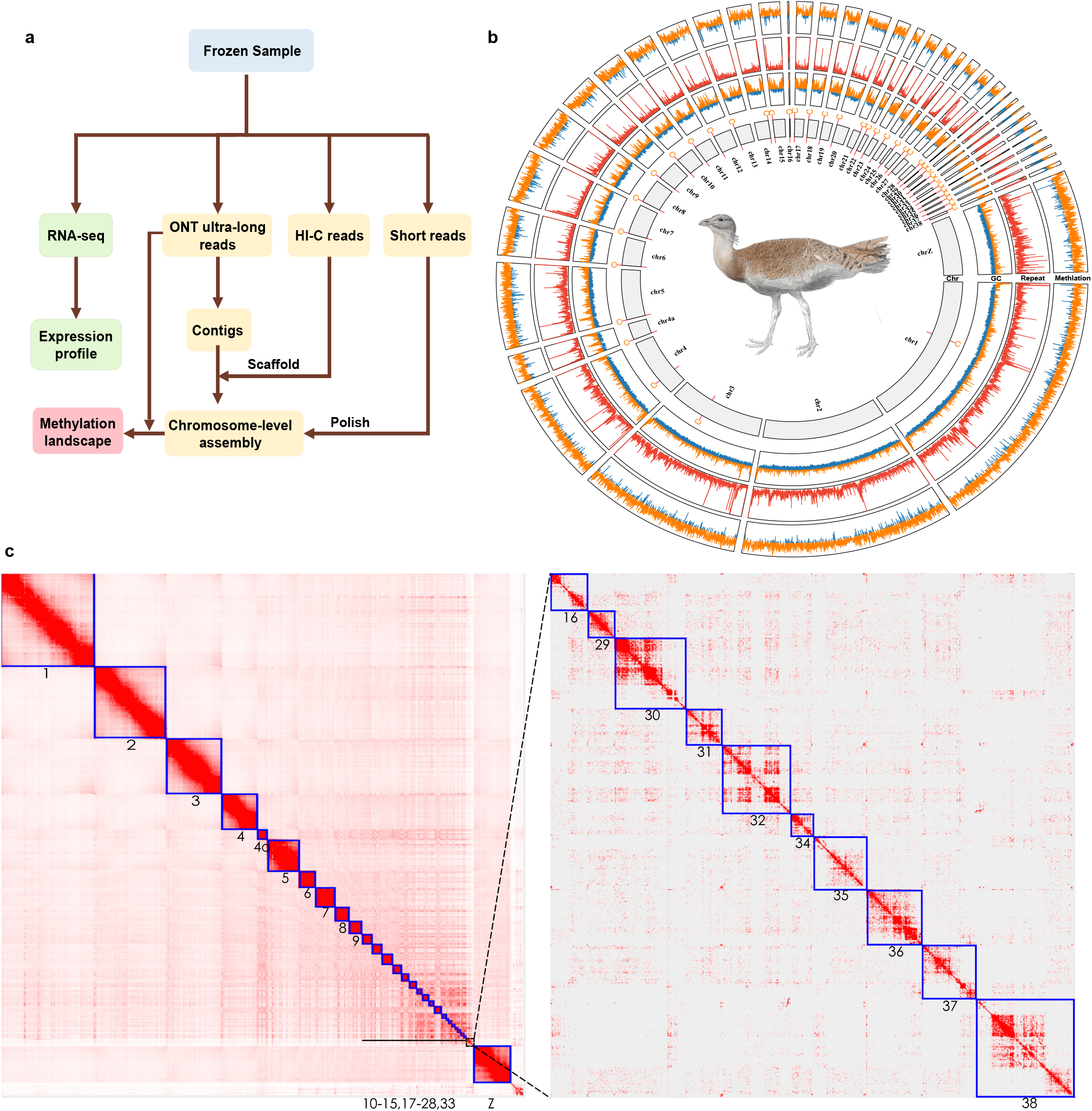
A near-complete chromosome-level genome assembly. **a**) a workflow of this study. **b**) Chromosomal sequence features of the great bustard. Chromosome IDs are labeled at bottom of chromosome bars. In the “Chr” ring, the “C” label represents the centromere position in each chromosome. In the “GC” ring, the orange bars represent regions (in 50 kb windows) with GC-content higher than the genome average (42.94 %), while the blue bars represent the below-average regions. In the “Repeat” ring, bars represent repeat content in 50Kb window. In the “Methylation” ring, the orange and blue bars represent regions with CpG-level higher or lower than 0.5, respectively **c**) Hi-C contact heatmap for the whole genome assembly visualized in Juicebox. The chromosome IDs are shown under the chromosome models. The right panel showed the zoom-in view for dot chromosomes (Chr16, 29-32, 34-38).

We further anchored the contigs to chromosome models using 58.6 Gb Hi-C data, generating the final assembly OTswu. The Hi-C heatmap revealed 40 chromosome models (**Fig. 1b**), consistent with the known karyotype (2n = 80) (Nishida C, 1981). Those 40 chromosomes include 39 autosomes and a Z chromosome, accounting for 97.7% of the assembled sequence. Out of the 40 assembled chromosome models, 32 contain zero gaps; the other eight chromosomes have only 18 gaps (**Supplementary Fig. S2**).

To further evaluate the completeness of chromosomal assembly, we searched for the presence of the telomere repeats (TTAGGG)n and centromeric sequence. We found that the telomeric repeats were present at the ends of 20 chromosomes with a mean length of 2.3 kb (**Supplementary Table S3**). We identified a putative 191 bp tandem repeat (Cen191) that was present in 38 out of 40 chromosomes. The Cen191 repeats appear at one end in all microchromosomes (**Fig. 1c, Supplementary Fig. S2-S3**), consistent with the acrocentric morphology of bird microchromosomes (J. Li et al., 2021).

The repetitive sequences occupy 15.1% of the great bustard genome (**Supplementary Table S4**), in contrast to the mean value of ~9.5% in other birds (Feng et al., 2020). This partially explains the larger genome size (~1.20G) of great bustard compared with the average bird genome size (~1.10 Gb) (Kapusta, Suh, & Feschotte, 2017). We annotated 19,898 protein-coding genes, with 17,512 (88%) of them functionally annotated for protein domains.

### 3.2 Extremely conserved karyotype throughout avian evolution

The diploid number of chromosomes (2n = 80) in great bustard is equal to that in emu which is thought to represent the ancestral avian karyotype (J. Liu et al., 2021). In contrast to the complete assembly of chromosomal models in OTswu, a few small microchromosomes are unfortunately absent in the emu genome assembly. We thus chose to compare OTswu to a new zebra finch genome assembly (bTaeGut1.4.pri) (J. Kim et al., 2022), and a recently released chicken genome (GGswu) that has complete chromosome models (Zhen Huang et al., Submitted). Our synteny analysis showed that all great bustard chromosomes have one-to-one homology with chicken chromosomes except for chr4 and chr4a (**Fig. 2a**) which were known to have fused in chicken (Itoh & Arnold, 2005; Völker et al., 2010). Compared with chicken or great bustard, the zebra finch genome experienced one fission leading to chr1 and chr1a (Itoh & Arnold, 2005) and one newly identified fusion between two small microchromosomes (dot chromosomes) (**Fig. 2a**). According to such chromosome comparisons, we inferred that the great bustard likely retains the ancestral avian karyotype, reflecting extreme conservation of karyotype during avian diversification (Ellegren, 2010; Z. Huang & Xu, 2022).

**Fig 2.**
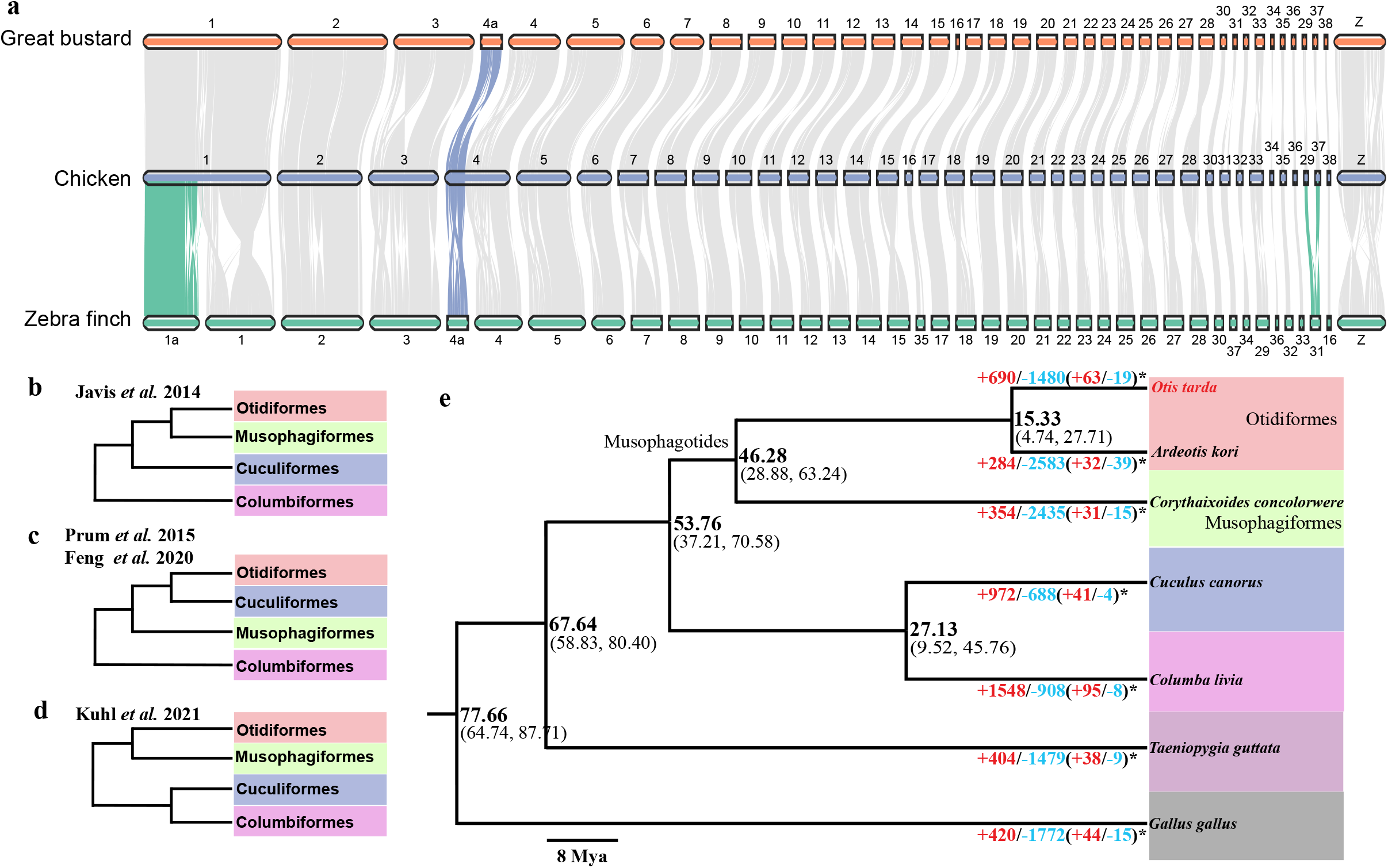
Chromosome evolution and phylogenomics. **a**) Gene synteny blocks among the great bustard chicken and zebra finch were identified and visualized by MCScan. Horizontal bars represent chromosome models. The synteny blocks highlighted in colors indicate chromosomal fusions. **b-d**) Previously proposed phylogenetic relationships for four bird orders. **e**) The ultrametric tree reconstructed by this study. Divergence time and confidence interval (95% HPD) are labeled at the nodes. The numbers below branches show the numbers of expanded (blue) and contracted (red) gene families. Numbers of rapid expanded and contracted gene families are labeled with *.

### 3.3 Phylogenomic

The phylogenetic position of Otidiformes is one of the unresolved problems in the bird tree of life despite large-scale phylogenomic efforts in the past decade. Using whole-genome alignment data, Jarvis *et al*. (2014) placed Musophagiformes as the sister group of Otidiformes (Jarvis et al., 2014) (**Fig. 2b**), while targeted capture data (Prum et al., 2015) suggested Cuculiformes to be the sister group (**Fig. 2c**). Though the Musophagotides clade (Musophagiformes+Otidiformes) is further supported by a transcriptome-based phylogeny (**Fig. 2d**) (Kuhl et al., 2021), a more recent phylogenomic study (Feng et al., 2020) again suggested that Cuculiformes is more close to Otidiformes. To construct phylogenetic relationships, we used whole-genome alignments and aligned coding sequences of 5,832 single-copy orthologous genes from seven birds. Those species include two Otidiformes, one Musophagiformes, one Cuculiformes, one Columbiformes species, and two outgroup species (chicken and zebra finch) (**Methods and Materials**). The phylogeny derived from whole-genome alignment (**Fig. 2e**) or coding sequence alignment was in agreement with Jarvis *et al*. (2014), though protein sequences-based phylogeny was identical to that of Kulh *et al*. (2021). Based on the topology we obtained with the whole-genome alignment, we estimated the divergence time across the phylogeny. Our analysis showed that Otidiformes diverged from Musophagiformes approximately 46.3 million years ago (**Fig. 2e**).

### 3.4 Methylation landscape correlated with GC content and gene expression

Across the great bustard genome 85.5% of the CpG sites are methylated in the muscle tissue, a percentage somewhat higher than that in human tissues (70-80%) (Suzuki & Bird, 2008). This percentage varies across chromosomes, with smaller chromosomes having much fewer methylated CpG sites (**Fig. 3a**). Notably, dot chromosomes have only 68.91% of the CpG sites methylated (**Fig. 3a**), though they contain a higher density of CpG sites and have higher GC content (**Fig. 1b**) than microchromosomes or macrochromosomes (**Supplementary Table S5-S7**). A lower percentage of methylated CpG sites at least in part explains lower chromosome-wide methylation (5mC) levels in dot chromosomes (0.501) than in microchromosomes (0.580) or macrochromosomes (0.602) (**Fig. 3b**, **Supplementary Table S8**). This is in contrast to a previous study in chicken where dot chromosomes were found to have higher methylation levels (Zhen Huang et al., Submitted). In chicken dot chromosomes the higher methylation levels are likely driven by the large hypermethylated heterochromatic regions which are, unfortunately, only partially assembled in OTswu (**Fig. 3c**). The dot chromosomes in the OTswu assembly instead contain mostly gene-rich euchromatic sequences (**Fig. 3c**).

**Fig 3.**
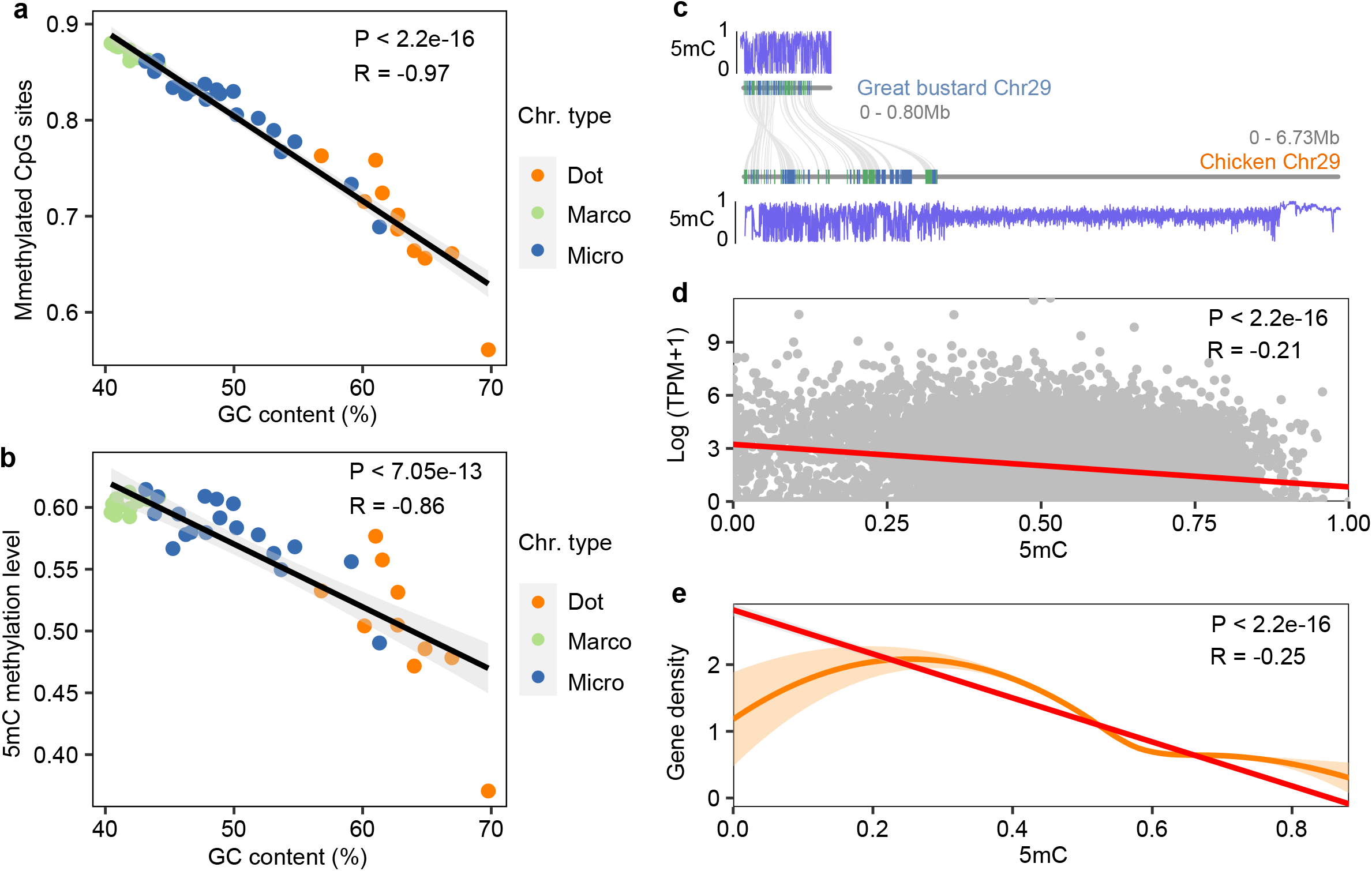
Methylation patterns across chromosome. **a**) The percentage of methylated CpG sites is negatively correlated with GC content. **b**) Chromosome-wide methylation levels show negative correlation with GC content. **c**) gene synteny between the chr29 of chicken and great bustard, and methylation landscapes of the two chromosomes. The chicken chr29, as one example of dot chromosomes, has a large hypermethylated gene-desert region. **d**) The expression levels of genes are negatively correlated with the gene-body methylation levels in muscles. **e**) The methylation levels are negatively correlated with gene density in 50 kb windows.

To investigate the relationships between DNA methylation and gene expression (Sepers et al., 2019), we collected RNA-seq data from muscles, the same tissue that we used to call methylation. We found that the methylation level and gene expression levels were significantly negatively correlated (Pearson’s correlation P < 2e-16, R= −0.21) (**Fig. 3d**), corroborating the role the DNA methylation in repressing gene expression (Bird, 2002). Our analysis also revealed that the methylation level was negatively correlated with gene density (Pearson’s correlation P < 2e-16, R= −0.25) (**Fig. 3e**).

### 3.5 Adaptive to powered flight

The expansion and contraction of gene families play an essential role in phenotypic diversification and adaptation to the environment (Albalat & Canestro, 2016). We identified 63 rapidly expanded and 19 contracted gene families in the great bustard genome (**Supplementary table S9**). Among the 63 expanded gene families are those related to ATPase, short-chain dehydrogenases reductases, mitochondrial inner membrane protein and immunoglobulin. Out of the 24 hierarchy of enriched GO terms for the 432 genes from the expanded gene families, ten directly related to cardiac functions, including regulation of cardiac muscle contraction by calcium ion signaling (GO:0010882) and cardiac muscle contraction (GO:0060048) (**Supplementary Fig. S4**, **Supplementary sheet 1**). The increased copy number of genes involved in cardiac contraction, actin contraction and calcium ion signaling transduction may play a role in enhancing blood circulation and oxygen transport during the long-range flight of the great bustard (Wu et al., 2021). Enhancing blood circulation may also be responsible for accelerating heat generation to endure cold environments (Duchamp & Barre, 1993; Adan, Ardevol, Remesar, Alemany, & Fernandez-Lopez, 1994).

We subsequently test positive selection signals in the great bustard genome. After three rounds of robustness tests, we detected 763 genes with positive selection signals (**Supplementary sheet 2**). GO Enrichment analysis for the positively selected genes (PSGs) showed significantly enriched terms mostly related to metabolism processes, including protein lipidation (GO:0006497), monocarboxylic acid catabolic process (GO:0072329) and lipid catabolic process (GO:0016042) (**Fig. 4a**, **Supplementary sheet 3**). Those PSGs have higher (FDR <0.0001, **Supplementary Table S10**) expression levels than non-PGSs in all five tissues we sampled, and have the highest expression levels in muscles (**Fig. 4b**).

**Fig 4.**
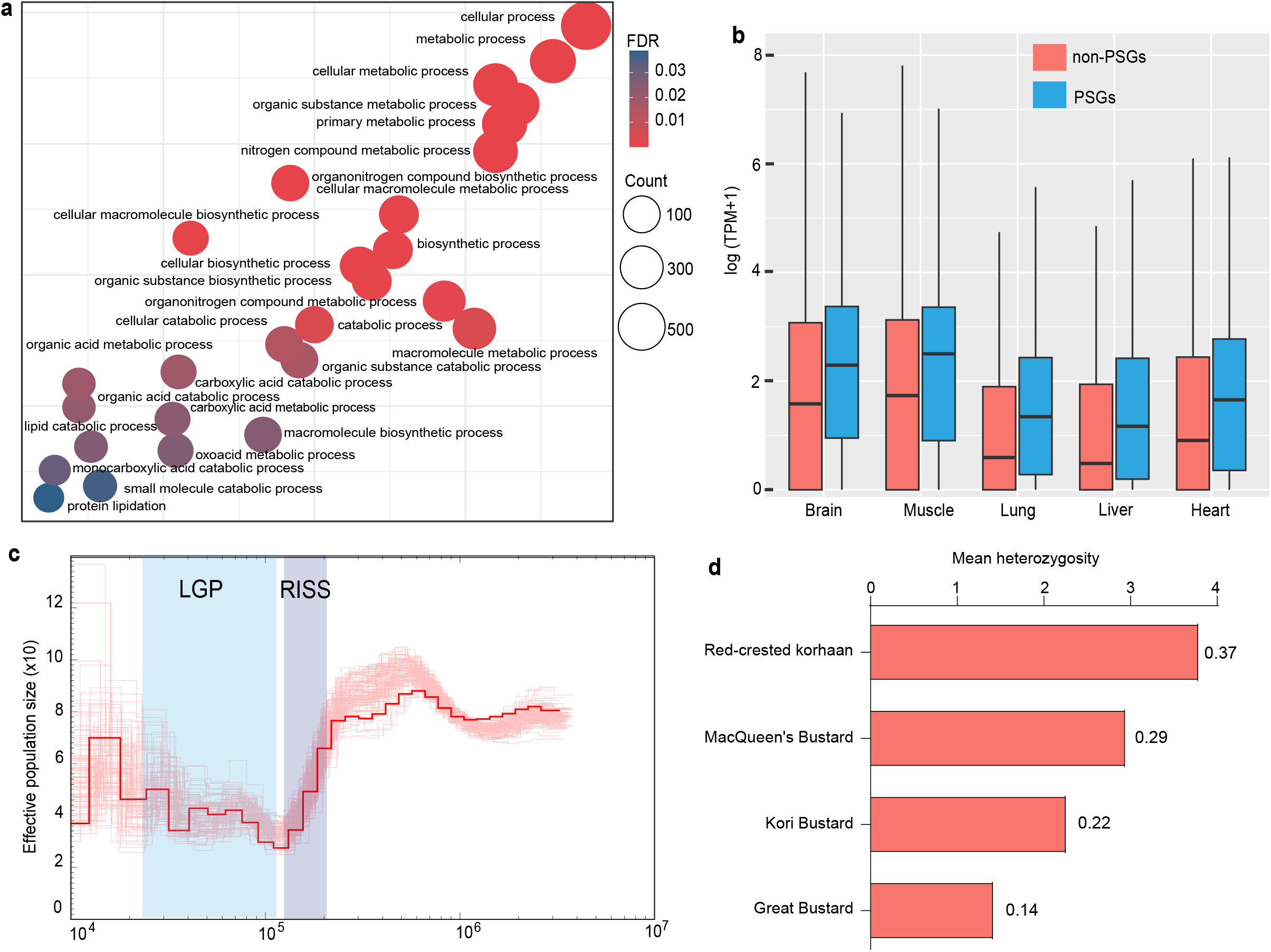
Adaptive evolution and demographic history. **a**). Enriched GO terms for PSGs. The sizes of filled circles represent counts of PSGs. The colors of circles indicate GO terms transpose FDR values. **b**) PSGs have higher expression levels relative to non-PGSs. **c**) The red curve is the PSMC based estimate for demographic history of the great bustard. The pink curves indicate 100 times of bootstrapping. **d**) The great bustard has lowest heterozygosity among four Otidiformes species.

### 3.6 Demographic history and genetic diversity

The genome-wide nucleotide divergence between great bustard and kori bustard was estimated to be 6.25%. We then estimated that the neutral mutation rate was 6.11e^10-9^ mutations per base per generation in great bustard, and used this value to infer population dynamics. The PSMC analysis suggests that the population effective size (*Ne*) was up to ~80,000 but decreased to ~30,000 from ~250 to ~20 Kya during the RISS glacial period. In the last glacial period (LGP), the effective population size stabilized, and gradually recovered to ~40,000. Since then, the effective population size fluctuated around ~40,000, similar to the census population size (**Fig. 4c**).

Due to the lack of population samples, we used individual genome heterozygosity to approximate the level of genetic diversity (Gelabert et al., 2020). The heterozygosity of great bustard was 0.14%, the lowest among the four Otidiformes genomes we sampled (**Fig. 4d**). MacQueen’s heterozygosity was twice larger than that of great bustard, consistent with the twice larger census population size (78,960 −90,000 Vs. 40,000).

### 3.7 A recently evolved sex chromosome stratum

Neoaves share a large non-recombining region of the ZW sex chromosomes, and many lineages further independently experienced an additional event of recombination suppression (Zhou et al., 2014; Kimball & Braun, 2022), leading to varying lengths of the pseudoautosomal regions (PARs). We demarcated the boundary of PAR in the great bustard by mapping female resequencing data to the male reference genome. A ~10 Mb sequence at the edge of the Z chromosome shows a diploid coverage and high female heterozygosity (**Fig. 5a-b**). However, if we used a stringent filtering criterion by removing the reads with mismatch numbers larger than 2, only a ~2 Mb sequence showed a diploid coverage (**Fig. 5c**). A likely explanation for the disparity is that the region from ~2 to ~10 Mb (S3) experienced a recent event of recombination suppression between the Z and W, and the W was insufficiently differentiated from the Z, allowing the W-derived reads to be mapped to the Z (Xu, Wa Sin, Grayson, Edwards, & Sackton, 2019). If this were the case, only the first ~2 Mb should be identified as the PAR.

**Fig 5.**
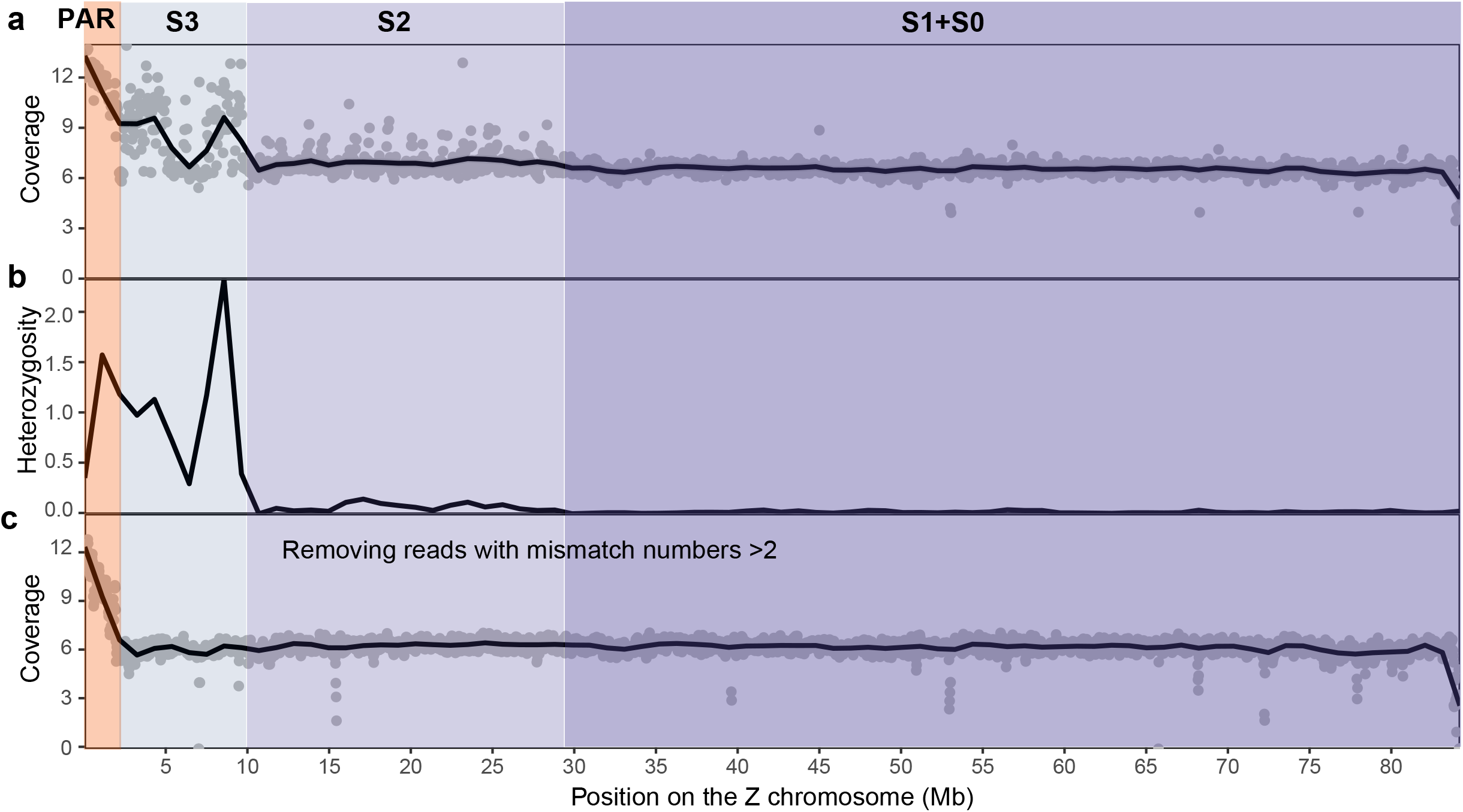
Identification of a young evolutionary stratum on the sex chromosome. **a**) The coverage of female re-sequencing data calculated in 50 kb windows. **b**) female heterozygosity calculated in 50 kn windows on the Z chromosome. **c**) similar to a) but the reads with mismatch number larger than 2 were removed from the alignments before calculating coverage.

Similarly, the region from ~10 to ~28 Mb showed a few heterozygous sites, in contrast to the rest of the Z being almost hemizygous (**Fig. 5b**), as well as fluctuation in coverage of alignments using standard mapping criteria (**Fig. 5a**). This region is likely the third evolutionary stratum (S2) whose range on the Z chromosome matched what has been identified on other Neoaves birds (Xu, Auer, et al., 2019). The boundary of S2 aligned with the breakpoint of an inversion between the Z chromosomes of great bustard and emu (**Supplementary Fig. S5**) that was predicted to create the Neognath S1 (Zhou et al., 2014).

## 4 Discussion

In this study, we generated a near-complete chromosome-level genome assembly of the great bustard using DNA extracted from a dead individual found in the wild. The OTswu ranks the fourth most continuous assembly among more than 800 bird genomes available in NCBI, in terms of contig N50. Moreover, all 40 chromosome models have been assembled. This has been rare in bird genome assemblies because the small microchromosomes, or dot chromosomes, are difficult to be assembled due to their high GC content and abundant tandem repeats (Peona et al., 2021; J. Kim et al., 2022). Our success in generating the near-complete genome assembly suggests that high-quality DNA were well-preserved in low temperature and likely a very recent death of the sampled individual. We have also extracted high-quality RNA which is known to degenerate much faster than DNA, allowing for transcriptome-based gene annotations and profiling gene expression across tissues. As the rate of DNA or RNA decay strongly depends on temperature and time since death (L. H. Huang et al., 2017), we recommend quick translocations of dead wild animals to the laboratory for DNA and RNA extractions. This may supply valuable samples for threatened or endangered species that are otherwise difficult to come by.

Great bustard has received increasing attention in recent years, and large efforts have been made in species conservation, including re-introduction (according to the Royal Society for the Protection of Birds). Our study provides a high-quality genome assembly of great bustard which will serve as an important genomic resource for conservation management. The low genetic diversity of great bustard revealed by our analysis is alarming, demanding future surveys of genetic diversity among great bustard populations.

Our comparative genomic analyses have also provided insights into the avian genome evolution. Because the sampled individual is male, the assembled genome lacks a female-limited W chromosome. Nevertheless, we were able to infer the evolutionary strata on the Z chromosome based on sequencing coverage and heterozygosity. Further efforts are needed to assemble a female genome in order to analyze the gene content and evolution of the W chromosome (Xu & Zhou, 2020), and similar analyses are required for closely related lineages, such as Musophagiformes, to illustrate a more complete evolutionary history of sex chromosomes. Our chromosomal comparison across Neognaths also provide direct genomic evidence for the evolutionary stasis of avian chromosome (Ellegren, 2010), even for the smallest microchromosomes that receive little study until recently (M. Li et al., 2022).

## Acknowledgments

We thank Junjie Yin (Xiamen University) for assistance in figure preparations. This study is supported by the start-ups funding from Southwest University to LX.

## Data Accessibility and Benefit-Sharing

The genome assembly data are available at GenBank under the accession number: JAPMTP000000000. Raw sequence reads are deposited in the BioProject PRJNA903785.

## Author Contributions

L.X., H.L., and X.O. conceived the project and designed the experiments. H.L., X.J, and J.W. and J.S. performed genome sequencing and assembly. H.L., X.J, and Z.X. performed data analyses. X.Z., Z.H., B.L, and M.H. participated in the project design and provided samples. H.L. and L.X. wrote the manuscript and revised the manuscript. All authors read and approved the final manuscript.

